# Targeting Integrated Stress Response by ISRIB combined with imatinib attenuates STAT5 signaling and eradicates therapy-resistant Chronic Myeloid Leukemia cells

**DOI:** 10.1101/2021.05.05.442756

**Authors:** Wioleta Dudka, Grażyna Hoser, Shamba S. Mondal, Laura Turos-Korgul, Julian Swatler, Monika Kusio-Kobiałka, Magdalena Wołczyk, Agata Klejman, Marta Brewińska-Olchowik, Agata Kominek, Milena Wiech, Marcin M Machnicki, Ilona Seferyńska, Tomasz Stokłosa, Katarzyna Piwocka

## Abstract

Integrated Stress Response (ISR) facilitates cellular adaptation to variable environmental conditions by reprogramming cellular response. Activation of ISR was reported in neurological disorders and solid tumours, but its function in hematological malignancies remains largely unknown. Previously we showed that ISR is activated in chronic myeloid leukemia (CML) CD34+ cells, and its activity correlates with disease progression and imatinib resistance. Here we demonstrate that inhibition of ISR by small molecule ISRIB, but not by PERK inhibitor GSK2656157, restores sensitivity to imatinib and eliminates CM Blast Crisis (BC) D34+ resistant cells. We found that in Patient Derived Xenograft (PDX) mouse model bearing CD34+ imatinib/dasatinib-resistant CML blasts with *PTPN11* gain-of-function mutation, combination of imatinib and ISRIB decreases leukemia engraftment. Furthermore, genes related to SGK3, RAS/RAF/MAPK, JAK2 and IFNγ pathways were downregulated upon combined treatment. Remarkably, we confirmed that ISRIB and imatinib combination decreases STAT5 phosphorylation and inhibits expression of STAT5-target genes responsible for proliferation, viability and stress response. Thus, our data point to a substantial effect of imatinib and ISRIB combination, that results in transcriptomic deregulation and eradication of imatinib-resistant cells. Our findings suggest such drug combination might improve therapeutic outcome of TKI-resistant leukemia patients exhibiting constitutive STAT5 activation.

## Introduction

Chronic myeloid leukemia (CML) which is driven by oncogenic BCR-ABL1 tyrosine kinase, is an example of a disease that is successfully treated with molecular targeted therapy. Introduction of imatinib significantly improved CML treatment, patients’ life expectancy and overall survival ^1,2^. However, although imatinib shows remarkable clinical efficacy in the chronic phase, the effects in advanced phases are short-lived, complete remissions are rare, and relapse occurs often ^3–5^. Many patients show primary or secondary resistance to imatinib or second generation tyrosine kinase inhibitors (TKIs), such as dasatinib, nilotinib or bosutinib. The resistance originates in majority from cellular intrinsic mechanisms. Apart from *BCR-ABL1* point mutations (e.g. T315I) which affect drug binding affinity ^6,7^, the *BCR-ABL1* gene amplification or clonal evolution may lead to relapse driven by both BCR-ABL1-dependent and -independent mechanisms.

The most recognized pathways responsible for resistance are mediated by activation of JAK2/STAT5, RAS/RAF/MAPK or PI3K/Akt/mTOR ^3,8,9^. They activate proliferation, anti-apoptotic response and survival, cytokine and growth factors signaling, altogether strongly promoting resistance to treatment and disease relapse. Therefore, targeting these pathways is one of the current strategies for eradication of resistant cells ^10–12^.

Previously, we identified that the PERK-eIF2α pathway related to Integrated Stress Response (ISR) is activated in CD34+ CML-BP cells ^13^. ISR is a highly conserved signaling responsible for cell adaptation and survival upon stress conditions ^14–17^. This is achieved by phosphorylation of the eukaryotic translation initiation factor eIF2α, remodelling of translation ^18^ and transcription of stress response effector genes, including CHOP and GADD34, which are ISR markers.

Under physiological conditions, the ISR is one of the mechanisms sustaining homeostatic balance in a healthy cell. Cancer cells can utilize ISR to survive and develop drug resistance. Previous reports demonstrated that ISR is active in solid tumors in which it correlates with hypoxia and metastasis ^19^. However, ISR has not been deeply studied in leukemia. Since recognized, ISR is proposed as a therapeutic target in cancer ^20–22^. Nevertheless, no efficient and specific strategy has been proposed still, especially for hematological malignancies.

We report here that inhibition of ISR signaling by small molecule ISRIB combined with imatinib has potential to eradicate imatinib-resistant CML-BP cells. We show that such treatment specifically changes gene expression profile and inhibits oncogenic STAT5 signaling. Therefore the combination of ISRIB and imatinib was identified as a possible therapeutic strategy when aiming to eradicate TKI-resistant leukemic cells exhibiting constitutive STAT5 activation.

## Methods

### Cell culture

The K562 cells (CCL-243) and LAMA84 cells (CRL-3347) were purchased from American Type Culture Collection (ATCC) and cultured ^13^. Cells were authenticated at ATCC service and were regularly tested for Mycoplasma contamination. Detailed description of the two-step generation of cells expressing non-phosphorylable form of eIF2α is provided in the Supplementary Information.

### Isolation of CD34+ CML-BP patient cells

CML CD34+ cells were obtained from the Institute of Hematology and Blood Transfusion in Warsaw, Poland, in accordance with the Declaration of Helsinki and with patients’ consent and approval of the local Ethical Committee (Ethical and Bioethical Committee UKSW, Approval No.WAW2/059/2019 and WAW2/51/2016, Approval No. KEiB-19/2017). The characteristics of patient is detailed in the Supplementary Information. Peripheral blood mononuclear cells (PBMC) were isolated by density gradient centrifugation and CD34+ cells were separated using EasySep human CD34+ selection cocktail (StemCell Technologies, Inc.). CD34+ cells were short-term cultured in IMDM medium (Invitrogen) with 10% FBS, 1 ng/ml of granulocyte-macrophage colony-stimulating factor (GM-CSF), 1 ng/ml of stem cell factor (SCF), 2 ng/ml of interleukin-3 (IL-3). Cells were cryopreserved and kept in −180C until usage.

### Cell treatment

Thapsigargin (Sigma) was used at 100 nM; imatinib (gift from Lukasiewicz Pharmaceutical Institute, Warsaw) at 0,5 or 1 μM concentrations *in vitro* or at given doses *in vivo*. ISRIB (Merck, SML0843) was given as indicated. GSK2656157 (GSK157) (Calbiochem) for *in vitro* test was dissolved in DMSO and given as indicated. For *in vivo* studies, first the step general stock of GSK157 was made (53,3 g of GSK157 to 1523 μl DMSO). In the second step 20 μl of the general stock of GSK157 was added to 44 μl of PEG400 (MERC, #8074851000) and 40 μl of saline (not PBS).

### *In vivo* experiments

Experiments were performed using immunodeficient NOD.Cg-PrkdcscidIl2rgtm1WjL/SzJ mice, in accordance with the Animal Protection Act in Poland (Directive 2010/63/EU) and approved by the Second Local Ethics Committee (Permission No. WAW/51/2016). Cells (10^6^) were injected subcutaneously or into tail vain. Mice were treated with: imatinib - twice a day (50 mg/kg); GSK157 - once a day (20 mg/kg); ISRIB - once a day (2 mg/kg) or in combination with the same doses, as indicated. Experimental schemes are presented as part of Figures.

### Flow cytometry

Apoptotic cell death was detected using Annexin V-PE Apoptosis Detection Kit I (BD Biosciences #559763) as described ^13^. To detect phosphorylation of STAT5 and S6K, cells were incubated with eBioscience™ Fixable Viability Dye eFluor™ 455UV (Thermo Fisher) to discriminate dead cells, followed by staining using Transcription Factor Phospho Buffer Set (BD Pharmingen) and antibodies: anti-phospho-STAT5 (Tyr694)-PE, and anti-phospho-S6 (Ser235, Ser236) – eFluor450 (eBioscience, Thermo Fisher). Events were acquired using BD LSR Fortessa cytometer (Becton Dickinson) and then analysed by FloJo Software (Becton Dickinson).

### Western Blot

Western blot analysis was performed in a standard conditions, as previously described ^13^. List of antibodies is presented in the Supplementary Information.

### RT-qPCR analysis

Total RNA was extracted using TRI Reagent (Sigma #T9424) or by Renozol (Genoplast #BMGPB1100-2) followed by Total RNA Mini column purification kit (A&A Biotechnology #031-100). 2 µg of RNA was subjected to reverse transcription using M-MLV enzyme (Promega #M1705), dNTP mix 100 mM each (BLIRT #RP65) and oligo (dT)_18_ primers (Bioline #BIO-38029). The RT-qPCR reaction was performed using SensiFAST SYBR Hi-ROX Kit (Bioline #BIO-92020) on the StepOnePlus™ platform (Thermo Fisher Scientific) according to MIQE guideline. Primers sequences are listed in the Supplementary Information. The comparative 2^-ΔΔCt^ method was used to determine the relative mRNA level using StepOnePlus software. 18SrRNA was used as a reference control. Data are presented as mean values ± SD; n = 3-5). Statistical significance was assessed using unpaired Student’s t-test with Welch’s correction and p ≤ 0,05 was estimated as significant (*p ≤ 0.05; **p ≤ 0.005; ***p ≤ 0.001; ****p ≤ 0.0005).

### RNA Sequencing and data analysis

RNA was isolated as described in RT-qPCR section. The library was prepared using NEB Next Ultra II Directional RNA library Prep kit for Illumina (#E7335S/L). Sample analysis: the quality of raw data was verified in FASTQ format from RNA-Seq experiments with FastQC ^23^. Because of observed high quality of the raw data, no further processing of reads was performed. Data analysis was done using the SquIRE ^24^ pipeline. Human genome hg38 and corresponding refseq gene annotations were downloaded from UCSC (https://genome.ucsc.edu/; ^25^ with SQuIRE. STAR version 2.5.3a ^26^, StringTie version 1.3.3b ^27^, and DESeq2 version 1.16.1 ^28^ were used within the SQuIRE pipeline for alignment of reads, transcript assembly and quantification, and differential gene expression analysis, respectively. Differentially expressed genes with false discovery rate (FDR) < 0.05 were reported here. Principal component analysis of all samples (11 replicates in total from 4 conditions) based on gene expression data (transcripts per kilobase million or TPM) was performed with ^29^. The Clust tool ^30^ was used for co-expressed gene clusters identification across all samples. The default normalization procedure of Clust for RNA-seq TPM data (quantile normalization followed by log2-transformation and Z-score normalization, code “101 3 4”) was applied. gProfiler ^31^ was utilized for the simultaneous functional enrichment analysis of the genes from all clusters in multi-query mode. The RNA-Seq data from this publication have been deposited to the NCBI GEO repository (https://www.ncbi.nlm.nih.gov/geo) and can be accessed with the dataset identifier GSE171853.

### Statistical analysis

Data were analysed using GraphPad Prism (GraphPad Software, La Jolla, CA, USA) Single comparisons were tested using unpaired Student’s *t*-tests for normal distributed samples or Mann–Whitney-U tests when normal distribution was not given. One-way or two-way ANOVA was applied for multiple comparison analysis, with Bonferroni’s multiple comparison post-test. For RT-qPCR unpaired Student’s t-test with Welch’s correction was applied. P values < 0.05 were estimated as significant (*p<0.05; **p <0.005; ***p<0.0005). Data are presented as mean ± SD.

## Results

### GENETIC ISR INHIBITION SENSITIZES CML CELLS TO IMATINIB *IN VITRO*

To study the impact of Integrated Stress Response globally, ISR was inhibited by targeting the main regulatory hub - eIF2α. This was achieved by expression of non-phosphorylable (S51A) eIF2α form (visible as additional band on western blot), followed by overexpression of shRNA against eIF2α 3’UTR (S51A shUTR) to inhibit expression of endogenous wt eIF2α, leading altogether to complete lack of eIF2α phosphorylation (Fig. 1A, detailed procedure of generation of genetically modified cells is provided in Supplementary Information). Both generated cell lines had unaffected levels of PERK, an UPR kinase acting upstream of eIF2α, and expressed GFP necessary for FACS sorting (Fig. 1A). The functional influence on ISR confirmed by detection of mRNAs encoding ISR markers *CHOP* and *GADD34* showed that inhibition of the eIF2α phosphorylation attenuates dynamics of the ISR activation (Fig. S1A). In addition, inhibition of the eIF2α phosphorylation itself decreased cell viability (Fig. 1B), and associated with increased basal *GADD34* and *CHOP* mRNA levels indicating stress-induced cell death (Fig. S1B). This indicated that the lack of ISR pathway itself is cytotoxic for CML cells. Furthermore, imatinib-induced apoptosis was higher in S51A and further increased in S51A shUTR cells, compared to wt (Fig. 1C). This implies that indeed K562 cells utilize the eIF2α phosphorylation-dependent mechanism and that ISR inhibition sensitizes CML cells to imatinib.

**Figure 1.**
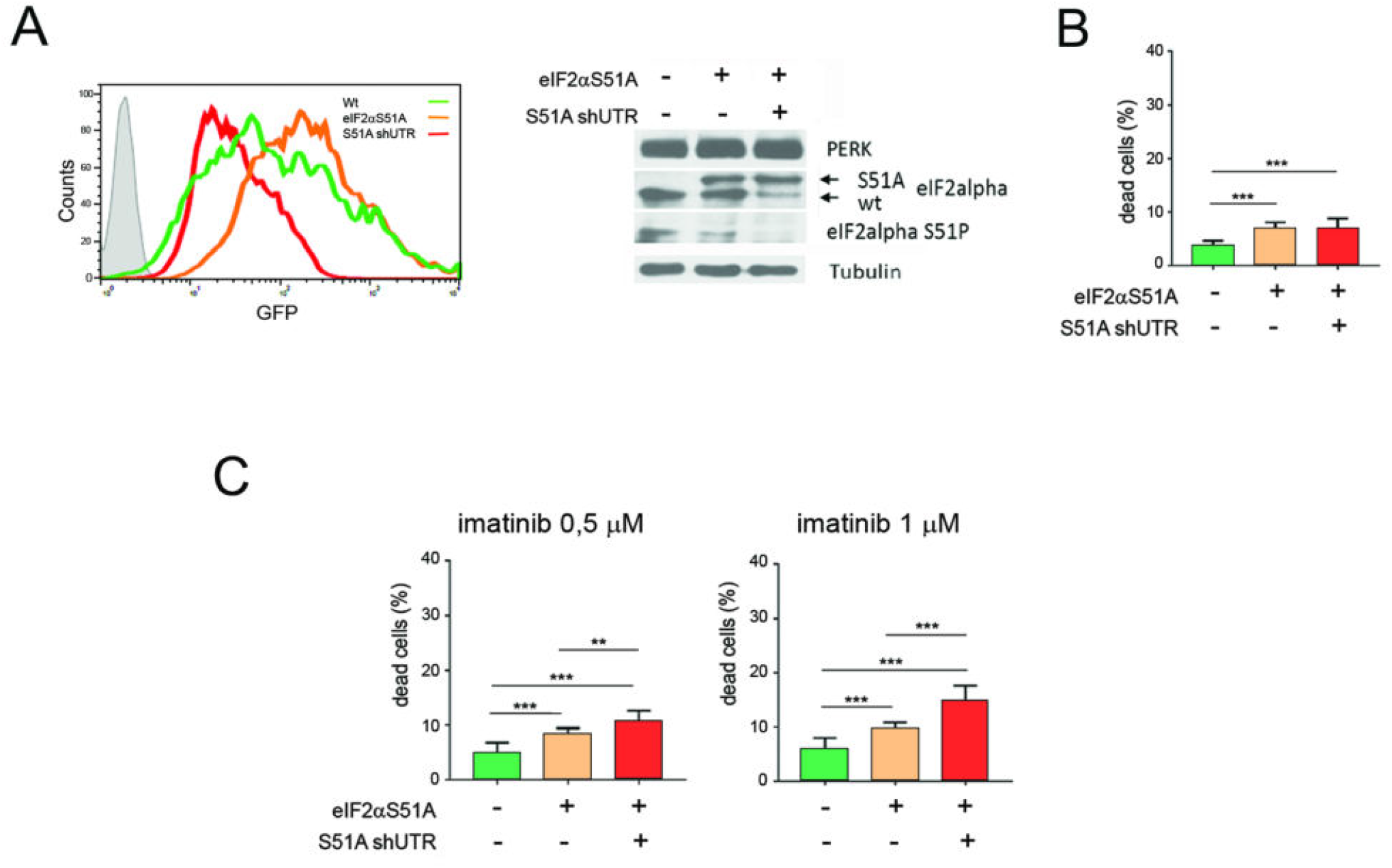
Genetic ISR inhibition by targeting eIF2α phosphorylation increases apoptosis induction and sensitizes K562 CML cells to imatinib *in vitro*. A. Left panel: transfection levels estimated by GFP fluorescence detection by flow cytometry. Overlay of representative histograms of K562 CML cells expressing wt eIF2α (green line), mutated non-phosphorylable form eIF2α S51A (orange line) and mutated form together with construct containing shRNA sequence against 3’UTR region of eIF2α (S51A shUTR – red line); Right panel: The levels of PERK, eIF2α and phosphorylated eIF2α (S51P) protein estimated by western blot in wt, or stably transfected eIF2α S51A and S51A shUTR mutants. Arrows indicate wt (lower) and mutated (40 kDa higher) eIF2α bands. Tubulin was used as a loading control. B,C. Cell death detected by flow cytometry in K562 wt, eIF2α S51A and S51A shUTR mutant cells in untreated conditions (B) or after treatment with 0,5 and 1 μM imatinib (C). Data are shown as a percentage of dead cells measured using AnnV/7AAD assay. Statistical analysis: Unpaired t test with Welch’s correction (*p ≤ 0.05; **p ≤ 0.005; *** p ≤ 0.0005).

### ISRIB, BUT NOT GSK157, SENSITIZES CML CELLS TO IMATINIB *IN VIVO*

Even if the genetic approaches are useful, the pharmacological inhibition still gives the highest possibility for the clinical applications targeting the signaling pathways. Thus, we tested two ISR inhibitors: GSK2656157 (GSK157) and ISRIB (Fig. 2A). GSK157 is an ATP-competitive inhibitor of PERK kinase, which stops the PERK-dependent ISR activation. Small molecule ISRIB blocks the eIF2α -P-dependent downstream signaling and inhibits the executive part of ISR, without the cytotoxic effects ^32–35^. Both drugs have not been tested in leukemia, including CML. Pre-conditioning of K562 CML cells with either GSK157 or ISRIB, followed by ISR induction by thapsigargin, revealed that both ISR inhibitors significantly reduced expression of *CHOP* and *GADD34* mRNAs in leukemia cells (Fig. 2B, C).

**Figure 2.**
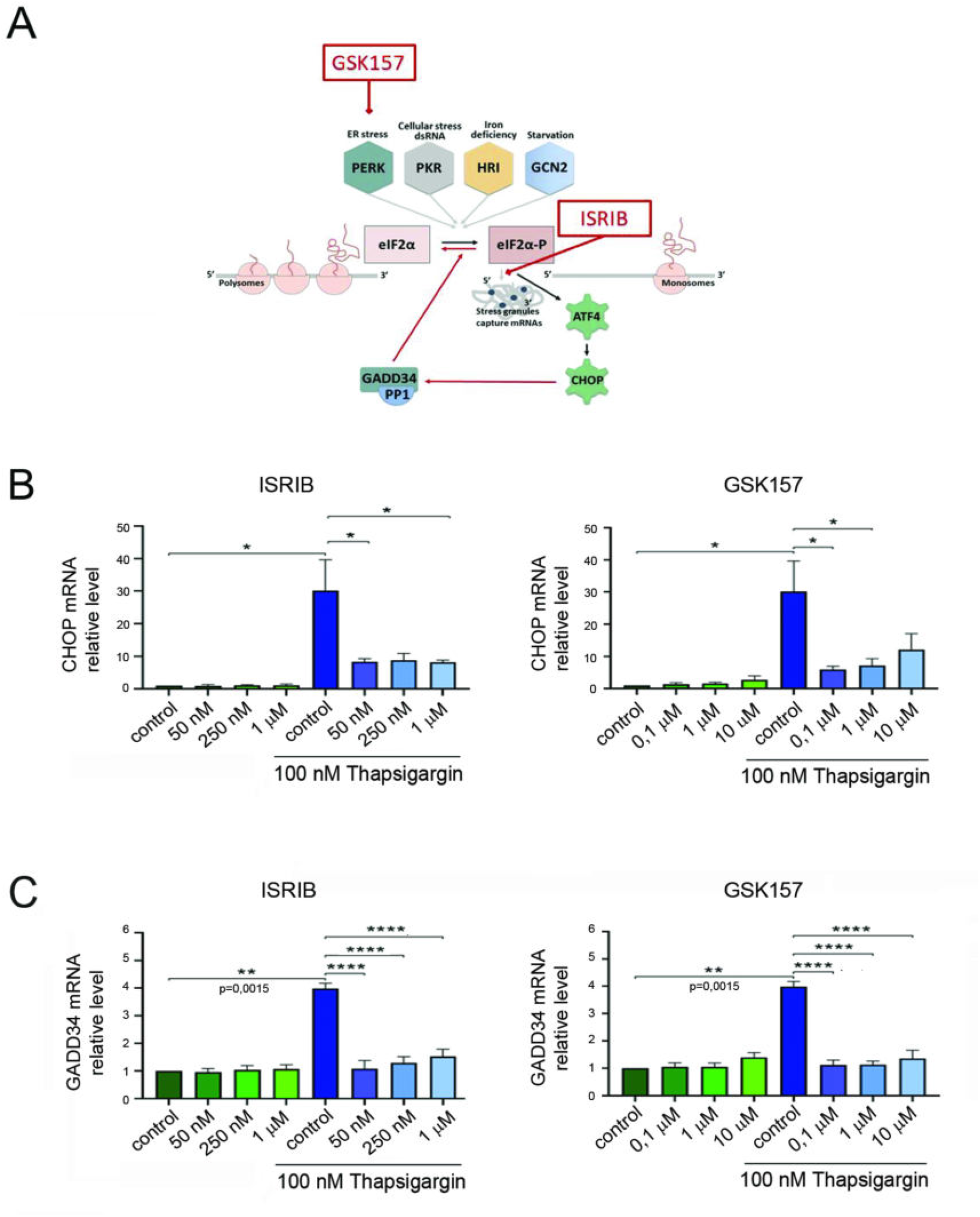
Pharmacological treatment with ISRIB or GSK157 impairs the ISR activation in K562 leukemic cells *in vitro*. A. Schematic graph of the ISR signaling pathway with the site of ISRIB and GSK157 action. B, C. CHOP and GADD34 mRNA expression levels measured by RT-qPCR in K562 cells. Cells were preconditioned with either ISRIB or GSK157 inhibitors in indicated concentrations, followed by ISR induction by 100nM thapsigargin for 2 hours. The level of not treated cells (CTR) was used as a reference =1. Statistical analysis: unpaired Student’s t-test with Welch’s correction and p ≤ 0,05 was estimated as significant (*p ≤ 0.05; **p ≤ 0.005; ***p ≤ 0.001; ****p ≤ 0.0005).

The results obtained *in vitro* (Fig. 2) imply that ISR inhibitors might improve the imatinib efficacy and eliminate CML cells. To test this hypothesis and verify cell survival and growth potential *in vivo*, xenograft studies were performed using NSG immunodeficient mice and GFP+ K562 cells (experimental scheme and treatment - Fig. 3A). After the time period necessary for leukemia cells growth, tumors were found in all untreated mice. Remarkably, only combination of imatinib and ISRIB significantly decreased number of animals with tumors (41% of mice developed tumors) (Fig. 3B). This was associated with the decreased tumor mass by more than 70% (Fig. 3C, D). Conversely, imatinib or imatinib with GSK157 exerted only moderate inhibitory effect, and the average tumour mass was not significantly different between those conditions. These results show that ISRIB but not GSK157 sensitizes CML to imatinib *in vivo*.

**Figure 3.**
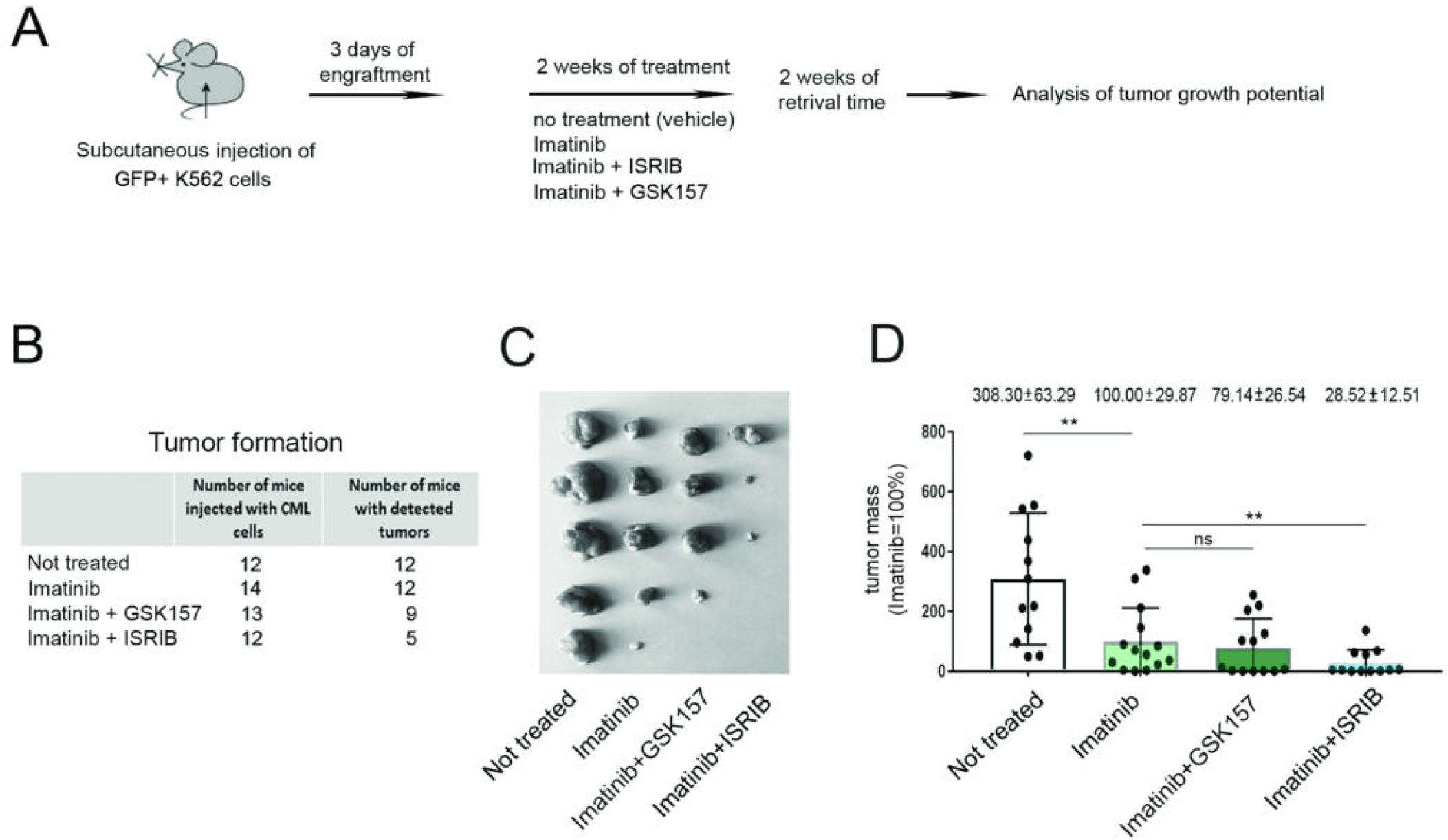
ISRIB, but not GSK157, sensitizes K562 CML cells to imatinib *in vivo*. A. The workflow of the *in vivo* experiment. Mice were: not treated (n=12); or treated with: imatinib (n=14); imatinib and GSK157 (n=13); imatinib and ISRIB (n=12). B. The number of mice which were injected with K562 cells and mice which developed tumors upon all tested conditions. C. The pictures of tumors isolated form representative experiment presenting the differences in proliferation potential of K562 cell in indicated variants. D. Corresponding graph showing the tumor mass. Tumors grown in mice injected with K562 cells and treated with imatinib were used as a control = 100%. Statistical analysis: Unpaired t-test, F-test to compare variances (*p ≤ 0.05; **p ≤ 0.005).

### ISRIB COMBINED WITH IMATINIB ATTENUATES ENGRAFTMENT OF PRIMARY TKI-REFRACTORY CML CD34+ BLASTS

Since the results in Fig. 3 implicated that combination of ISRIB and imatinib might eradicate imatinib-resistant CML cells and decrease the leukemia development, it was of paramount importance to verify this in the PDX model using NSG mice as a host bearing CD34+ CML cells resistant to imatinib and dasatinib. CD34+ cells were isolated from patient diagnosed in Blast Phase (BP), who initially responded to high dose imatinib, but subsequently developed resistance to imatinib, despite lack of detected (at the time of resistance) mutations within the kinase domain of BCR-ABL1. Dasatinib was introduced, but yielded only a transient effect. Next-generation sequencing revealed pathogenic variant in *PTPN11* gene described in hematological malignancies ^36,37^. The detailed patient characteristics are provided in the Supplementary Information. *PTPN11* gain-of-function mutations result in overactivation of the RAS/RAF/MAPK/ERK and the JAK/STAT pathways, in addition to their possible activation caused by BCR-ABL1. Thus, such cells represent a BCR-ABL1-independent, imatinib/TKI resistant phenotype.

A short and aggressive 7-day regimen was applied to test the beneficial effects, followed by treatment with imatinib or ISRIB alone or with drug combination (experimental scheme - Fig. 4A). All variants showed noticeable but not significant decrease in the spleen weight (Fig. 4B). To estimate the short-term engraftment, the percentage of human CD45+ (hCD45+) was detected within the bone marrow or spleen populations (Fig. 4C, 4D, 4E; Fig. S2). Combination of imatinib and ISRIB significantly decreased percentage of hCD45+ cells in the bone marrow, showing a 2-to 3-fold lower level compared to the treatment with imatinib or ISRIB alone. In addition, the combined treatment treatment decreased the engraftment into the spleen (which is considered as a secondary niche), compared to imatinib alone (Fig. 4E). These results showed that the combined treatment eradicates resistant blasts and decreases leukemia engraftment, therefore confirming the synergistic effect of imatinib and ISRIB.

**Figure 4.**
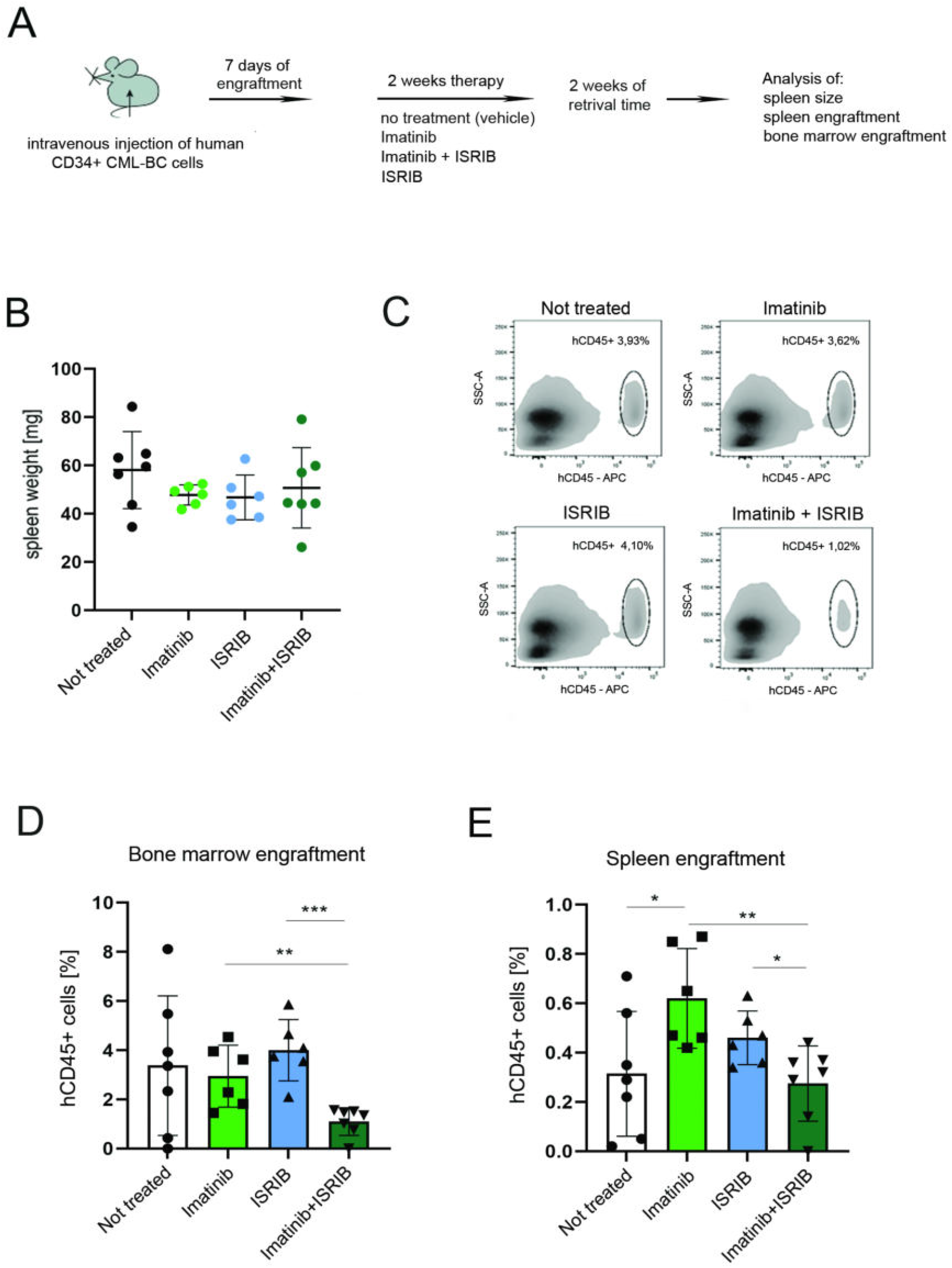
ISRIB in combination with imatinib attenuates engraftment of primary TKI refractory CML CD34+ blasts. A. The workflow of the *in vivo* experiment. PDX mice were: not treated/vehicle administrated (n=7); or treated with: imatinib (n=6); ISRIB (n=7); or combination of imatinib and ISRIB (n=7). B. Weight of spleens isolated from mice not treated or treated as indicated. C. Representative density plots showing the engraftment of hCD45+ CML primary cells into the bone marrow population under therapeutic treatment, detected by flow cytometry. hCD45+ population is gated on the hCD45 vs SSC dot plots, the percentage of hCD45+ cells is indicated. D, E. Corresponding graphs showing the bone marrow (D) or spleen (E) engraftment estimated by flow cytometric detection of hCD45+ CML primary cells in bone marrow or spleen, respectively, in given variants of treatment. The percentage of hCD45+ cells is shown. Statistical analysis: Unpaired t test, F test to compare variances (*p ≤ 0.05; **p ≤ 0.005; ***p ≤ 0.0005).

### COMBINATION OF ISRIB AND IMATINIB REPROGRAMMES THE GENE EXPRESSION PROFILE OF PRIMARY TKI-RESISTANT BLASTS

To investigate the molecular effects of the double treatment, RNA-seq was performed on FACS-sorted hCD45+ CML cells isolated from the PDX bone marrow. Principal component analysis (PCA) indicated that cells treated with imatinib and ISRIB are transcriptionally distinct (Fig. 5A). This was confirmed by hierarchical clustering of significantly changed genes between pairs of tested conditions (treatment vs control), and supported by the Pearson correlation values, which showed higher correlation between sole ISRIB and sole imatinib treatment compared to control (r = 0.69), than between each of the single treatments and the combined imatinib+ISRIB treatment compared to control (r = 0.32 and 0.37, respectively; Fig. 5B). The *SGK3* and *SNURF/SNRPN* genes regulating alternative RNA processing were identified as significantly downregulated upon the double treatment. Upregulated genes in majority encoded proteins regulating transcription and RNA processing.

**Figure 5.**
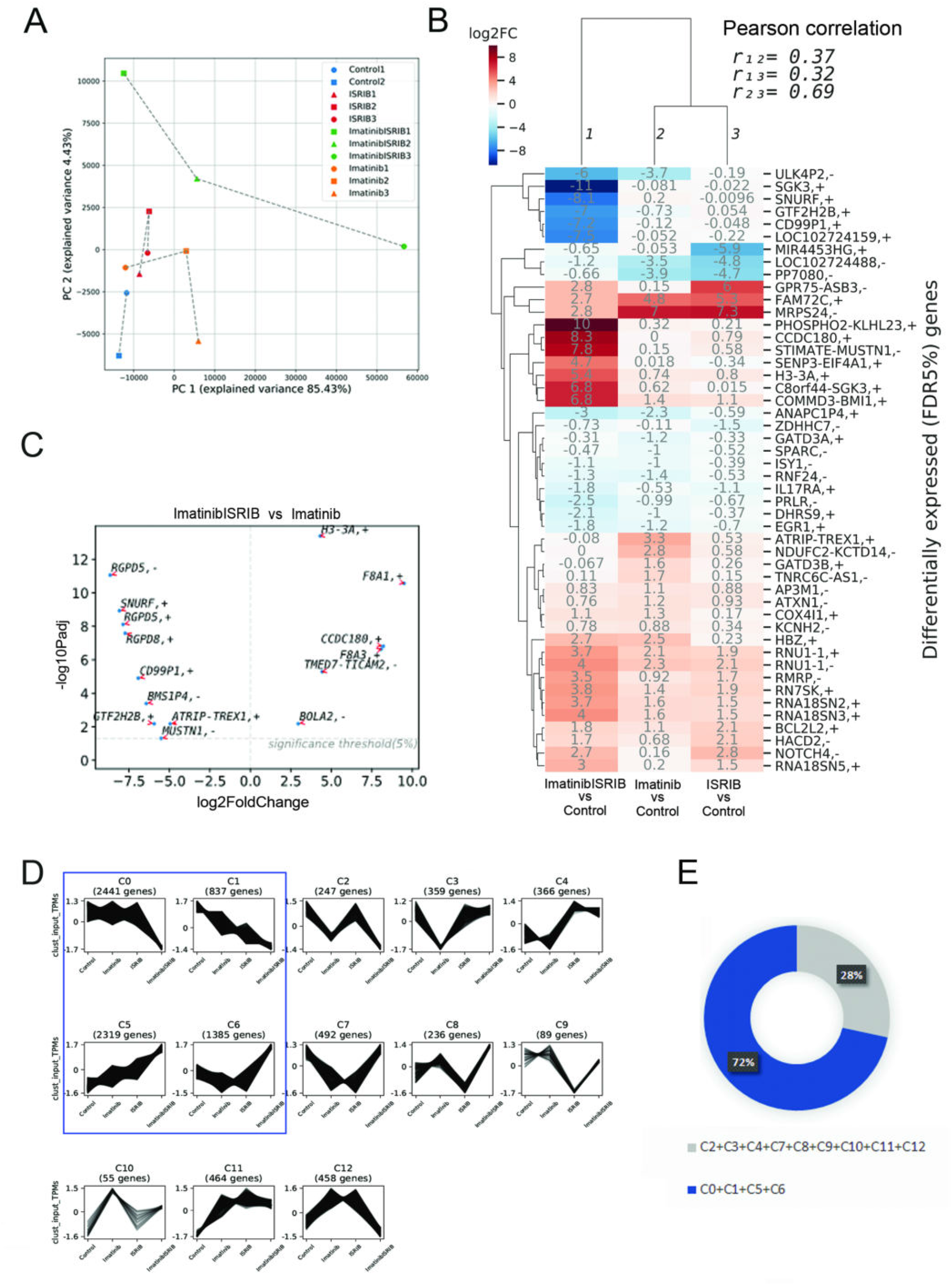
Combination of ISRIB and imatinib results in reprogramming of gene expression profile of primary TKI-resistant blasts. A. Two-dimensional principal component analysis plot of samples based on gene expression (TPM) data obtained from FACS-sorted hCD45+ CML cells isolated from untreated control mice (n=2, blue), or treated with ISRIB (n=3, red), imatinib alone (n=3, orange) or with combination of imatinib and ISRIB (n=3, green). B. Hierarchically clustered heatmap of fold-changes in expression (log2FoldChange) of significantly differentially expressed genes between the indicated pairs of conditions. Pairwise correlations of expression fold-changes are also shown. C. Significantly altered genes upregulated (positive value on x-axis) or downregulated (negative value on x-axis) in combined imatinib and ISRIB treatment versus with imatinib alone. D. Clusters (C0-C12) of co-expressed genes with varying patterns of gene expressions across all variants of treatment. Clusters C0, C1 displaying sharp downregulation or C5, C6 showing sharp upregulation of gene expression after combined treatment are marked in blue frame. E. Diagram showing the percentage of genes identified in four selected clusters C0, C1, C5, C6 (blue) and the rest (grey).

To identify genes responsible for the increased sensitivity, the gene expression profiles for imatinib versus imatinib + ISRIB were compared. In addition to the previously described (Fig. 5B), genes encoding proteins from the small GTP-binding RAS superfamily (*RGPD5* and *RGPD8*) were significantly downregulated (Fig. 5C, for all treatment combinations see Fig. S3A).

Since genes that are co-expressed are often co-regulated, clusters of co-expressed genes (C0-C12) across all variants of treatment were identified (all genes included, regardless of their statistical significance of change in expression) (Fig. 5D). Clusters with the highest number of genes represented the groups in which drug combination led to either sharp decrease (C0, C1) or increase (C5, C6) of gene expression (Fig. 5D, 5E). Those four clusters represented about 72% of all detected genes (Fig. 5E). This clearly indicated that the gene expression pattern for ISRIB + imatinib combination is specific and different from the other treatment conditions.

### COMBINATION OF IMATINIB AND ISRIB DOWNREGULATES GENES RELATED TO PROLEUKEMIC SIGNALING

To predict the cellular mechanisms altered by combined treatment, all 13 defined gene clusters underwent the functional enrichment analysis. The C0 and C1 clusters which included genes downregulated upon combined treatment, were significantly enriched in terms related to RAS/RAF/BRAF/MAPK signalling (Fig. 6, marked in red, and Fig S5; for all clusters see Fig. S3). Specifically, for Ras and MAPK signaling (detailed member genes in Fig. S4A, S4B), genes encoding RAF1, ARAF, ERK2, KRAS, SRC, JAK2 and a number of proteins involved in activation of MAPK cascade such as: MEK1, MAPK1, MAP4K1, MAP3K3, MADD were downregulated upon combined treatment. The drug combination attenuated also IFNγ signalling and immune response, which in leukemia can additionally mediate activation of the JAK2/STAT5 pathway and inflammatory response (Fig. 6, marked in blue; for all clusters see Fig. S3B). Downregulation of processes essential for leukemia-promoting kinase-dependent signaling and immune response was also confirmed by Gene Ontology Biological Processes (BP) terms (Fig. S5, see C0 and C1 clusters).

**Figure 6.**
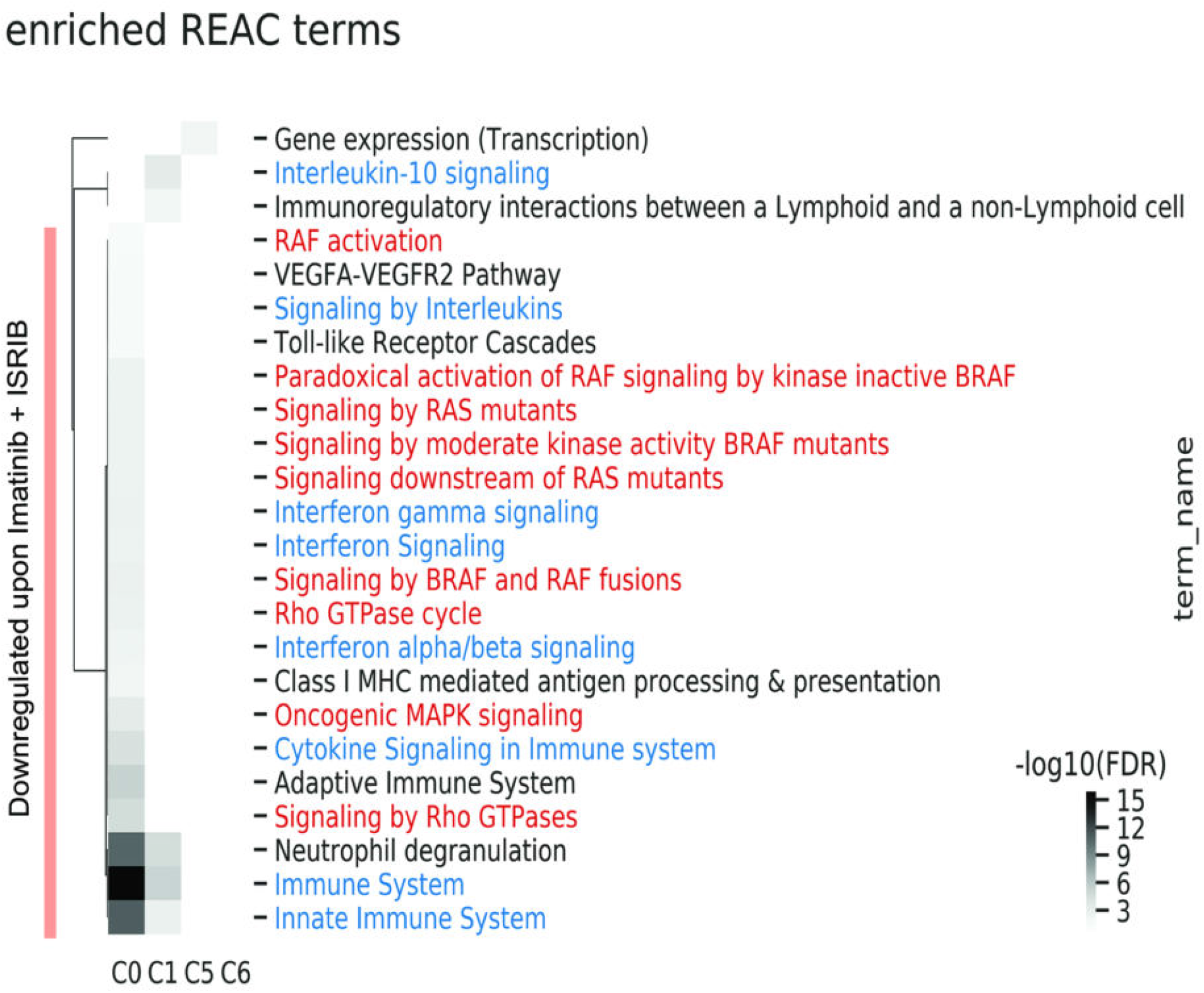
Co-expressed genes downregulated upon combined treatment are related to RAS/RAF/BRAF and Interferon gamma signaling. Functional enrichment Reactome (REAC ^66^) terms significantly enriched in C0, C1, C5 and C6 clusters. Downregulated genes belonging to C0, C1 clusters are indicated. RAS signaling is marked in red color, Interferon gamma signaling is marked in blue color.

While *SGK3* gene encoding serine/threonine-protein kinase SGK3 was significantly downregulated after the combined treatment (Fig. 5B), expression of the SGK3 interaction partners, selected based on the interaction partner datasource: BioGRID, IntAct (EMBL-EBI) and APID databases (see Supplementary Information), showed rather moderate inhibition upon combined treatment (Fig. S6). Among the downregulated genes, we found GSK3β, what may suggest its regulatory connection with SGK3 and specific downregulation upon combined treatment.

Altogether, these results showed that genes related to oncogenic pro-leukemic signaling were downregulated upon combination of imatinib and ISRIB, presumably enhancing targeting of leukemia cells by imatinib.

### COMBINATION OF ISRIB AND IMATINIB INHIBITS STAT5 SIGNALING IN CML CELLS

Transcriptomic data indicated that the combined treatment can downregulate oncogenic RAS/RAF/MAPK, JAK2, SGK3 and IFNγ signalling. In addition, genes that are mediators of JAK2/STAT5 signalling were attenuated (Fig. 6, Fig. S4, S5). This indicated that the combined treatment might inhibit the STAT5 pathway.

To obtain direct evidence that treatment with imatinib + ISRIB shows synergistic effect, STAT5 phosphorylation was assessed in K562 and LAMA84 cell lines, which were shown to activate the above signaling pathways ^13,38^. To better visualize the effects, strong ISR response *in vitro* was activated by thapsigargin. In both cell lines, combination of ISRIB with imatinib decreased STAT5 phosphorylation detected by western blot (Fig. 7A, 7B), and confirmed by phospho-flow cytometry (Fig. 7C). On the other hand, ISRIB alone did not change, whereas imatinib alone only partially decreased phosphorylation of STAT5, compared to double treatment, with effectivity lower in K562 cells, which were more resistant. Conversely, the significant additive effect (estimated by the phosphorylation levels) of the combined therapy combined to imatinib alone was not observed for other pro-leukemic related regulators such as: AKT, mTOR, S6K, SGK3, GSK3β or ERK (Fig. S7). So, the genetic data indicating downregulation of the SGK3 - GSK3β link were not confirmed *in vitro*. Interestingly, inhibition of AKT and ERK phosphorylation by imatinib (Fig. S7A and S7F, respectively), associated with decreased BCR-ABL1 activity (Fig. S8A), but not BCR-ABL1 protein level (Fig. S8B), indicated that either pAKT or pERK are not involved in acquiring the BCR-ABL1-independent resistance.

**Figure 7.**
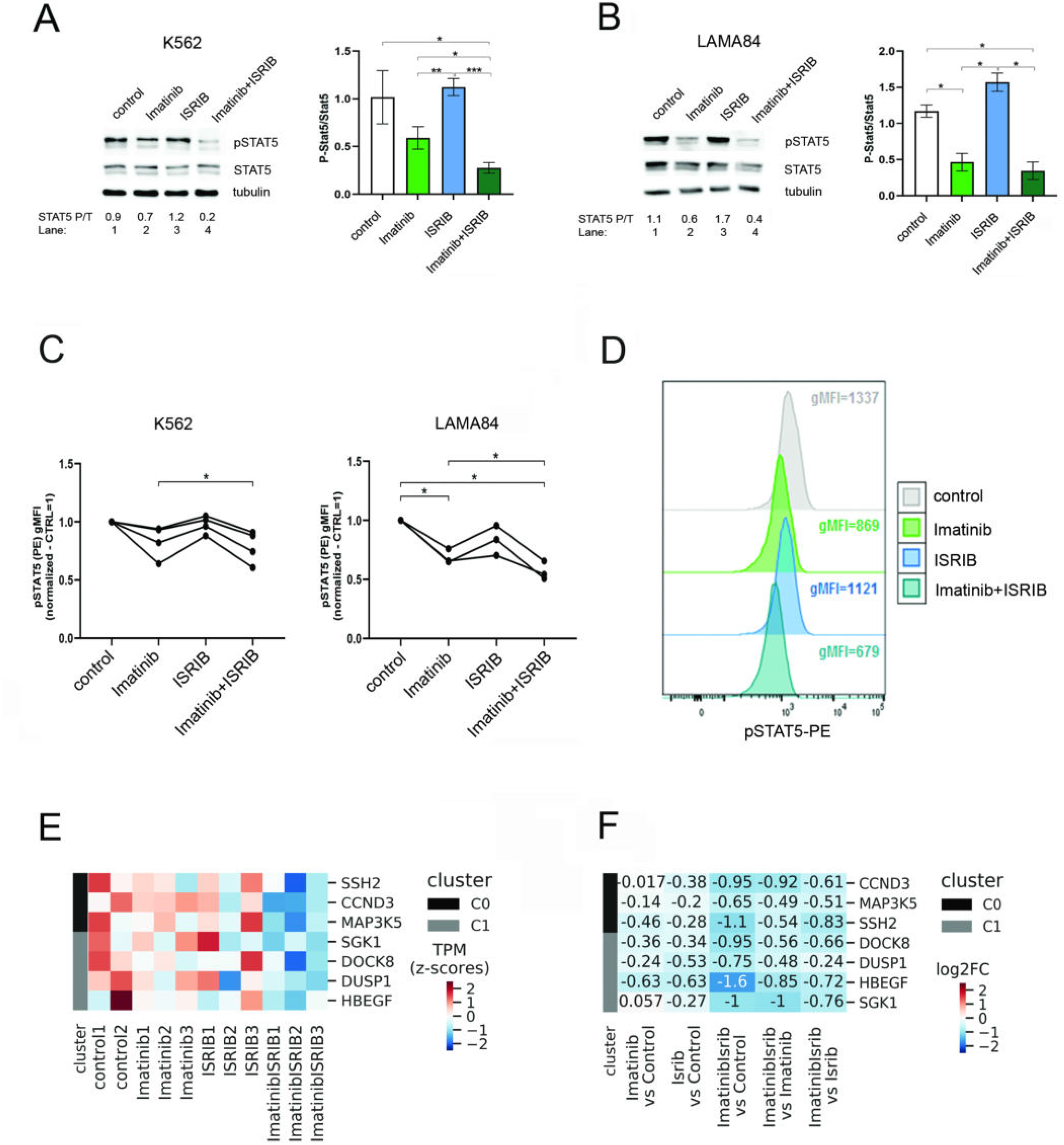
Combination of imatinib and ISRIB attenuates STAT5 signaling in CML cells. A, B. Left panels: Protein levels of STAT5 and phosphorylated form of STAT5 (pSTAT5) detected by western blot in K562 (A) or LAMA84 (B) CML cells untreated (control) or treated with drugs as indicated (all variants tapsigargin treated). The ratio of phosphorylated to total STAT5 forms (P/T) calculated based on the densitometry signal is given for each condition. A, B. Right panels: Adequate graphs showing pSTAT5/STAT5 ratios in K562 cells (A) and LAMA84 CML cells (B). Signal for control cells (without drug treatment) =1. Statistical analysis: unpaired T-test with Welch’s correction (*p ≤ 0.05; **p ≤ 0.005; ***p ≤ 0.0005). C. Flow cytometry analysis of pSTAT5 levels in K562 (left panel) and LAMA84 (right panel) cells untreated (control) or treated as indicated. Data were calculated based on gMFI, fluorescence signal for untreated cells=1. Statistical analysis: repeated-measures one-way ANOVA, with Tukey’s multiple comparisons test (*p ≤ 0.05). D. Overlay of the representative histograms presenting fluorescence signals for pSTAT5 estimated in control cells or in cells after treatment. gMFI values are indicated for each condition. E. The heat map showing expression level (transcript per kilobase million or TPM, standardized with z-score) of STAT5-target genes belonging to C0, C1 clusters shown for each gene across all replicates of untreated (control) and treatment conditions. F. The change in expression of STAT5-target genes belonging to cluster C0 and C1 in treatments comparison: expression fold change (log2FoldChange) in all comparisons.

The results presented in Fig. 7A-7D imply that the combined treatment attenuates the STAT5-dependent signaling. To test this, the fold change analysis of the STAT5 target genes expression was performed within the C0 and C1 clusters (downregulated upon imatinib + ISRIB). The list of possible STAT5 target genes was created based on ChIP-Seq data from malignant/hematopoietic cells (see Supplementary Information). As expected, the combined treatment decreased expression of STAT5 target genes (*SSH2, CCND3, MAP3K5, SGK1, DOCK8, DUSP1 and HBEGF*), compared to control or single treatments (Fig. 7E, 7F). Negatively regulated STAT5-target genes encoded regulators of cell cycle/proliferation, stress response and survival, including Slingshot Protein Phosphatase, Cyclin D3, ZIR8, MAP kinase phosphatase 1, EGF-like growth factor, MAP3K5 and SGK1. Data for all clusters are presented in Fig. S9. Conversely, such inhibitory effect was not observed for imatinib and ISRIB alone. Altogether, obtained data clearly support the conclusion that combination of imatinib and ISRIB shows the substantial synergistic effect and inhibits the proleukemic STAT5 signaling in CML-BC TKIs resistant cells.

## Discussion

Development of imatinib has revolutionised CML treatment and patients’ overall survival. Despite the clinical success of imatinib in the CML-CP treatment, the disease is still not fully curable and eradication of all leukemic cells is not efficient. Imatinib intolerance or primary resistance occurs, as well as many patients develop secondary resistance due to activation of signaling pathways, including JAK/STAT5, GSK3β or RAS/MEK/ERK ^3,8,9^. Importantly, such activation might occur in a BCR-ABL1-independent manner, thus upon imatinib treatment of even BCR-ABL1 non-mutated cells, those oncogenic pathways still remain active. Therefore, one of the current strategies to eradicate leukemic blasts, is to target BCR-ABL1 together with oncogenic signaling pathways, to resensitize cells to TKIs ^5,39–41^.

Here we provide evidence that inhibition of Integrated Stress Response by ISRIB combined together with imatinib might significantly break the resistance by targeting both, the stress response adaptative signaling as well as the STAT5-dependent intrinsic signaling. This can result in effective elimination of imatinib-refractory cells in CML. Unexpectedly, only ISRIB but not another ISR inhibitor - GSK157 belonging to the PERK inhibitors family, was effective *in vivo*. This is consistent with recent studies of amyotrophic lateral sclerosis which showed similar data indicating that ISRIB but not GSK157 inhibitor, was more effective and improved neuronal survival ^42^. Such effect can be a result of an eIF2α phosphorylation-independent effects ^43^, moderate specificity of GSK157, as its affinity to RIPK1 was shown to be significantly higher than to PERK kinase ^44^, as well as the pancreatic toxicity reported recently ^45^. Moreover, as PERK inhibitors target only one of four ISR arms, it can not be neglected that another parallel signaling leading to ISR is still active *in vivo*. In addition, ISRIB may have other, yet undescribed, targets. Results presented here provided several possible signaling pathways which may be altered by ISRIB in malignant cells.

ISRIB molecule, discovered in 2013, in contrast to PERK inhibitors, acts below eIF2α and directly reverses attenuation of the eIF2B by phosphorylated eIF2α ^33,46,47^. ISRIB has been proposed as a promising drug in the brain malignant conditions and age-related memory decline ^48,49^, as well as in some metastatic tumours ^50,51,52,53^. Recent studies showed that chemotherapy combined with ISRIB abrogates breast cancer plasticity and improves the therapeutic efficacy ^19^. This observation strongly supports the statement presented here. Studies of the clinical potential of ISRIB in hematological malignancies are limited ^54,55^. This study is the first to show ISRIB effectiveness in a combined therapy against CML-BP TKI-resistant blasts.

Mechanistically, we have discovered that the combined treatment inhibits STAT5 phosphorylation and decreases expression of STAT5 target genes, that regulate proliferation, apoptosis and stress response. Targeting STAT5, which is an oncogenic signaling in imatinib resistant forms of CML ^3,9,56^, effectively overcomes resistance and eradicates leukemic cells ^57–59^. The experimental therapy proposed by us, not only inhibits ISR but also attenuates the STAT5-dependent signaling in CML. It is to note, that the overactivated STAT5 has also been detected in other hematopoietic malignancies, such as non-CML chronic myeloproliferative disorders correlating with JAK2 V617F mutation ^60^ or Flt3-ITD positive AML ^61^. Therefore, it is worth considering that the proposed strategy might be effective also in other blood disorders.

In striking contrast, even if downregulation of related genes was observed in the transcriptomic analysis, the mTOR, SGK3, GSK3β, AKT and ERK activity was not specifically targeted by the double treatment *in vitro*. Notably, even though the regulatory ISR-SGK3 link was shown in glioma ^62^, and our transcriptomic data indicated SGK3 downregulation by the combined treatment, this was not confirmed in the model studies *in vitro*. On the other hand, pAKT and pERK, together with BCR-ABL1 activity were inhibited already by imatinib alone, and not further downregulated by drug combination. Therefore, those pathways were probably not responsible for the resistant phenotype. Nevertheless, in other leukemias in which the resistance developed due to AKT or ERK overactivation, such effect might help to eradicate the resistant blasts.

Interestingly, differential expression of genes responsible for the immune modulation (visible even in the xenograft model, which excludes involvement of T and B lymphocytes, but still encompasses functional myeloid cells) suggests possible involvement of the immune system remodelling in the therapeutic outcome. This data support the idea of targeting the innate immune system or immune checkpoints in myeloid malignancies, including CML ^63–65^. Thus, even though experiments were performed in immunodeficient (lacking adaptive, lymphocyte-mediated response) mice, signaling and functional effects related to the innate immune responses (mediated by e.g. macrophages) were possibly functional leading to the observed changes. Although interesting, this has to be verified in subsequent studies using the syngenic mouse model.

In conclusion, we discovered a novel strategy to break the resistance and eradicate imatinib-refractory CML blasts, which is based on therapeutic combination of ISR inhibitor ISRIB together with imatinib. We postulate that such strategy can improve therapeutic outcomes in CML patients showing TKI resistance related to overactivated STAT5 and stress adaptation signaling. Possibly, a similar approach based on ISRIB combined with a typical chemotherapy may also be applied to other hematological malignancies with constitutively activated STAT5 signaling and STAT5-dependent resistance.

## Acknowledgements

The research was funded by National Science Centre grants: PRELUDIUM (2015/19/N/NZ3/02267) to WD, Sonata Bis (2013/10/E/NZ3/00673) to KP and HARMONIA (2014/14/M/NZ5/00441) to TS. KP was also supported by TEAM-TECH Core Facility Plus (POIR.04.04.00-00-23C2/17-00) grant from Foundation for Polish Science co-financed by the European Union under the European Regional Development Fund. Authors would like to thank Dr. Antonis E. Koromilas for providing non-phosphorylable eIF2α S51A mutated form of eIF2α and valuable discussions, Lukasz Bugajski for help with part of experiments by FACS sorting, Danuta Wasilewska and Zuzanna Sipak for animal breeding and tissue isolation, and Jelena Pistolic and Vladimir Benes for help with RNA Seq library sample preparation. RNA Seq analyses were performed at the Gene Core EMBL Laboratory in Heidelberg.

